# Effects of Extruder Dynamics and Noise on Simulated Chromatin Contact Probability Curves

**DOI:** 10.1101/2025.06.09.658566

**Authors:** V. Konstantinov, A. Shadskiy, T. Lagunov

## Abstract

Loop extrusion by SMC complexes is a key mechanism underlying chromatin folding during both interphase and mitosis. Despite this shared mechanism, computational models of loop extrusion often rely on fundamentally different assumptions: interphase models typically use dynamic extruders with finite lifetimes, whereas mitotic models employ static extruders placed according to loop size distributions. In this work, we investigate whether these modeling paradigms are interchangeable or yield intrinsically incompatible results. Using publicly available Hi-C data from mitotic chicken cells, we systematically compare dynamic and static loop extrusion models implemented in the Polychrom framework. We evaluate how key parameters such as the extruder lifetime, extrusion velocity, and spatial noise affect the simulated contact probability curves P(s) and loop size distributions. Our results reveal that while both model types can be tuned to approximate the general shape of P(s), they produce distinct internal structures and divergent relationships between loop size and contact decay. We also show that increased extruder lifetimes lead to excessive nested loop formation, which alters both loop statistics and P(s) derivatives. Introducing spatial exclusion constraints between extruders partially restores consistency with static models. These findings highlight that differences in extruder behavior and polymer noise levels can significantly impact chromatin model outcomes and must be carefully accounted for when interpreting or comparing simulation results across biological conditions.

**Author summary:** Chromatin organization plays a crucial role in gene regulation and cellular function, yet our understanding of its three-dimensional structure relies heavily on computational modeling and the interpretation of complex experimental data. In this study, we use coarse-grained modeling approaches to simulate chromatin folding and systematically investigate how different analysis metrics and data processing methods influence the conclusions drawn from such models. By comparing widely used metrics and exploring the effects of normalization and noise, we highlight potential pitfalls and biases that can arise in chromatin modeling studies. Our findings provide practical recommendations for researchers in the field, aiming to improve the robustness and reproducibility of computational analyses of chromatin architecture. This work will help guide future studies toward more reliable interpretations of chromatin structure and its biological implications.

## Background

In recent years, a range of chromosome conformation capture techniques have been developed to study the spatial organization of the genome (1–5). Among them, Hi-C has emerged as one of the most informative and widely used methods, providing genome-wide maps of chromatin contacts and enabling detailed insights into three-dimensional chromosome architecture (3). However, Hi-C data are subject to inherent limitations, including restriction-enzyme-dependent biases arising from the choice of restriction enzyme (e.g., DpnII used in this study), and modest genomic resolution (∼1–10 kbp). These factors may influence the observed contact patterns and thus potentially affect model calibration and interpretation. While other methods such as Micro-C or approaches combined with ChIP-seq offer higher resolution and capture higher-order interactions, the broad availability and established analytic frameworks of Hi-C make it a reliable foundation for computational modeling (6–10).

One of the most prominent of these is the loop extrusion (LE) model, which posits that ring-like proteins of the structural maintenance of chromosomes (SMC) family assemble and load onto chromatin. Once loaded, these complexes actively translocate in opposite directions along the DNA fibre, extruding loops of chromatin in a directional manner (7,11). This leads to the emergence of loop organization in chromatin. Current evidence suggests that this mechanism plays a critical role in both interphase chromatin organization (7) and mitotic chromosome condensation (8,12).

While the core biochemical mechanism of loop extrusion is thought to be shared between interphase and mitosis (13,14), computational implementations of SMC activity in these two contexts often differ substantially. For interphase chromatin, a widely used model assumes dynamically loaded cohesin complexes that bind randomly, extrude loops by sliding their legs in opposite directions, and dissociate with a constant probability (7,15). These extruders are halted upon collision with neighboring complexes or oriented CTCF proteins, forming an ensemble of loops with stochastic sizes and limited lifetimes.

In contrast, mitotic chromosome models commonly adopt one of two approaches. A popular static framework posits that condensins are sequentially loaded onto chromatin, where they extrude loops until they are blocked by neighboring extruders, after which they become fixed. In this scenario, condensin II first establishes a dense array of long-range loops, followed by condensin I, which forms nested loops within this scaffold (8,12). This hierarchical assembly captures the experimentally observed loop sizes and density, and the model’s validity is supported by a good match between predicted and measured condensin occupancy per megabase (12). Furthermore, the rapid stabilization of loop sizes observed in the experimental data lends additional support to the static model. However, the static approach typically requires imposing artificial constraints, such as cylindrical confinement and helical axis generation, to recapitulate the global mitotic chromosome morphology (8,12). Other modeling strategies attempt to circumvent the need for cylindrical confinement by introducing additional physical features, such as increased interextruder affinity (16) or hypothetical “superextruders” (17) capable of generating compact mitotic structures through enhanced activity.

In contrast to the static approach, dynamic models of mitotic loop extrusion assume that extruders bind and unbind chromatin with a fixed probability, analogous to their behavior in interphase (11,18,19). Importantly, the residence time of these extruders is considerably shorter than the duration of mitosis. This framework reaches a dynamic equilibrium in which statistical properties—such as loop size distributions, the number of nested loops, and other structural metrics—stabilize over time. To reproduce mitotic chromosome patterns under this scheme, it is often assumed that extruders are capable of bypassing one another, forming so-called “z-loops” (18,19). This assumption is motivated by in vitro observations (20) and experiments involving bacterial condensins (21).

Despite this shared core mechanism, computational implementations of SMC complexes differ substantially between interphase and mitosis, reflecting distinct biological contexts and objectives. In interphase, dynamic loading and stochastic dissociation of cohesin are motivated by the need to simulate steady-state chromatin architecture that continuously remodels over hours. In contrast, mitotic chromosomes must rapidly condense and remain stable throughout mitosis (∼45 min in mammalian cells), suggesting a quasistable loop architecture established by early-loaded condensins that persist with minimal dynamics. Furthermore, in mitosis, the high density of condensin loading and tight coordination with topological constraints and chromosome geometry place different computational demands on the model. Static extruder frameworks, which capture the final-state loop architecture, often provide enough phenomenological descriptions of mitotic organization, even though the underlying SMC mechanism remains fundamentally dynamic at the molecular level.

In the present study, we sought to evaluate whether a dynamic extrusion model, operating without z-loop formation, is sufficient to recapitulate mitotic chromosome architecture under conditions of SMC3 and CAP-H2 subunit knockout. These specific experimental conditions were selected because of the reported absence of cylindrical chromosome morphology (12), thereby eliminating the need to impose external constraints in the simulation.

One commonly analysed feature is the contact probability as a function of genomic distance, P(s). This dependence is sensitive to both the polymer’s fractal dimensions—which characterize the self-similar, scale-invariant folding patterns inherent to polymer physics and the specific compaction regimes and loop architectures that emerge at different genomic scales depending on loop-extrusion parameters. These two factors interact: whereas an unconstrained polymer exhibits a universal fractal dimension (∼1.5 for a linear random coil), the introduction of loop extrusion modifies the effective fractal dimension at different scales, resulting in scale-dependent compaction patterns. Thus, analysing P(s) provides a sensitive readout of both fundamental polymer properties and extrusion-driven organization, making it a powerful but nontrivial method for model validation (8,9,18).

In this study, we used publicly available data and baseline parameters from (12) to assess how P(s) is influenced by the following:

- the choice of modeling framework (we reimplemented the model via the Polychrom toolkit as opposed to the original platform);
- the type of SMC model (dynamic vs static extrusion);
- the level of Gaussian noise added to the simulated contact maps.

Additionally, we compare several metrics for evaluating model-experiment agreement and demonstrate the strong dependence on the normalization point used in P(s) comparisons for most of them.

The focus of this study is a systematic evaluation of the reproducibility and comparability of chromatin modeling results under varying assumptions and implementations. Rather than limiting our investigation to a single mechanistic hypothesis, we conducted a multilevel comparison across commonly used modeling strategies.

First, we assessed the influence of different repulsive interaction potentials— specifically, the Step (22) and DPD (23) potentials—on the simulation outcomes. This comparison was performed using parameters closely aligned with those reported in prior studies to isolate the effect of the potential itself.

Second, we contrasted two fundamentally distinct extrusion models under matched conditions: a static model, in which loops are defined by a preassigned distribution, and a dynamic model, in which extruders stochastically bind and dissociate from chromatin.

Our results indicate that even when nearly equivalent input parameters are used, the choice of modeling framework and software implementation (in this case, the Polychrom toolkit versus the original platform) can substantially influence simulation outcomes.

This highlights the critical importance of careful validation and reporting of all the details of the computational model to ensure reproducibility in the field of computational modeling of chromosome architecture.

## Results

### Used simulations and experimental data

In this study, we utilized experimental Hi-C data generated using the DpnII restriction enzyme from mitotic chicken DT40 cell cultures, in which a degron system targets subunits of both the condensin II and cohesin complexes (12; see Methods). Under these conditions, the resulting contact probability curves P(s) lack statistically stable genomic contacts beyond 2 Mbp, as more distant interactions are presumably influenced by mechanisms unrelated to direct loop extrusion. As noted in the original study, such experimental perturbation is interpreted to limit the effective range of loop extrusion activity to approximately 2 Mbp.

Simulations were conducted via a modified version of the open2c/polychrom software package (22), which supports modeling chromosome architecture via dynamic loading and unloading of extruders. The extruder parameters were selected based on those reported in (12). As in that study, we did not use cylindrical confinement or include compartmentalization effects in our simulations. However, in contrast to that study, we did not impose any length constraints on the chromosomes, allowing the simulated chromosomes to reside in a diluted regime without external geometric restrictions.

### Baseline Outcomes Depend on Potential and Implementation

We began by evaluating the reproducibility of the experimental data via a model of a 40 Mbp chromosome with statically placed condensin I motors. The extruders were distributed according to an exponential loop size distribution with a mean of 100 kbp and an interextruder distance of 400 bp, parameters previously suggested as optimal for recapitulating contact density profiles in experiments (12). To assess the consistency of modeling results across different implementations, we employed two parameter sets and interaction potentials within the Polychrom framework: the Step potential, used as the default in dynamic extrusion models (24), and the DPD potential, utilized in (12,25). The Step potential characterizes beads as nearly ‘hard spheres’ with a steep potential gradient at the bead radius, minimizing overlaps outside the interaction zone. Conversely, the DPD potential models ‘softer’ beads with a smoothly distributed potential gradient over the interaction range. These two potentials are representative of typical ‘hard’ and ‘soft’ coarse-grained interactions, and are broadly comparable to other potentials used in polymer modeling. Further details are provided in the Methods section.

The resulting contact probability curves (Fig 1A) revealed discrepancies between the simulated contact distributions (as a function of genomic distance) and the experimental data for both potentials. These differences can be summarized as follows:

**Fig 1.**
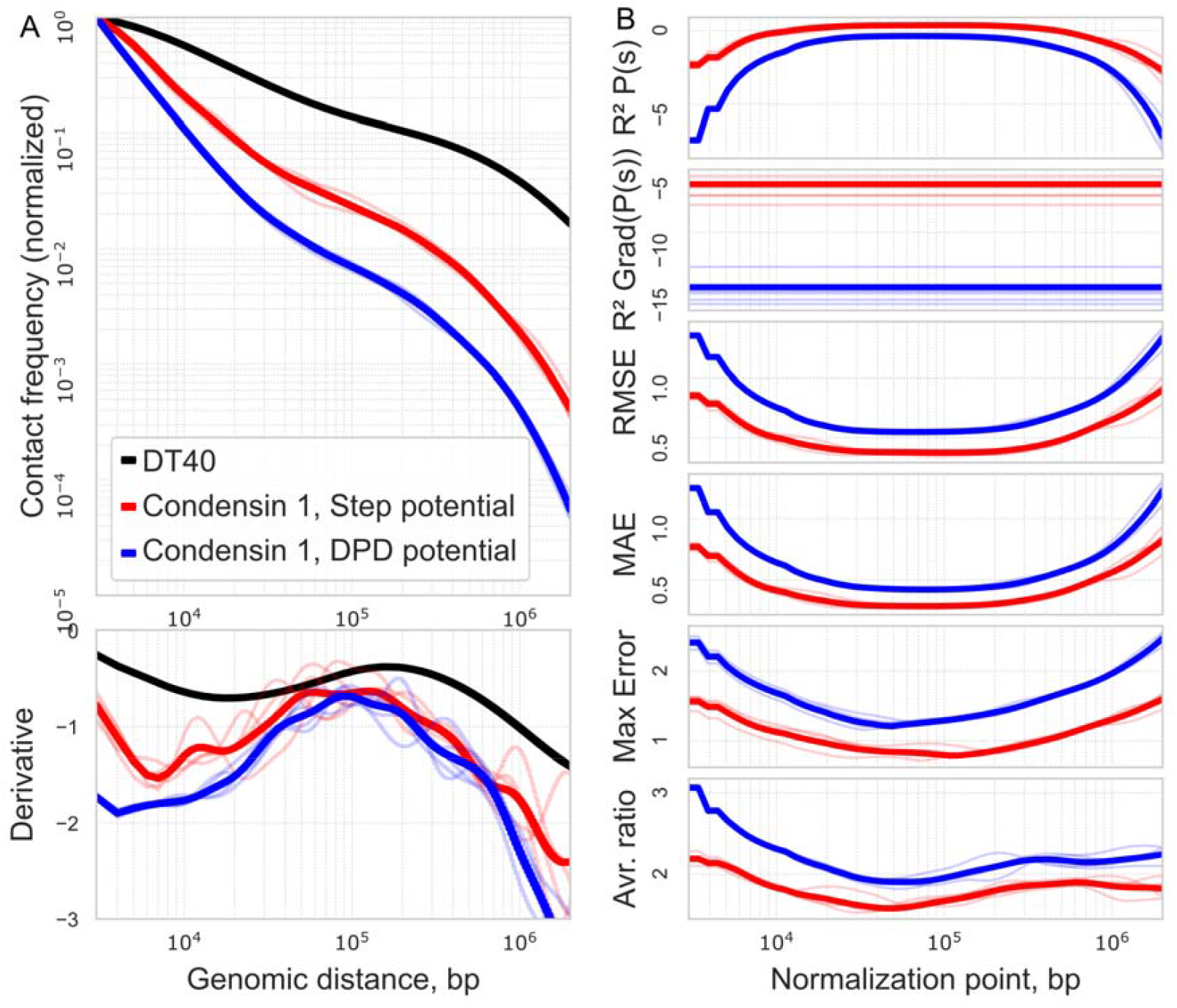
Comparison of experimental and simulated contact probability P(s) profiles. A) P(s) curves in the range of 3 kbp - 2 Mbp for experimental data for the chicken cell line DT40 [(12)] (black line), raw model single cells (transparent lines), aggregated raw model data (solid lines). Static extruders with exponential distribution of loop sizes: with Step potential (red line) and DPD potential (blue line); B) Results of comparing segments experiment P(s) vs model data with shifting normalisation points on different metrics. The normalisation point shifts within the range of 3 kbp to 2 Mbp. The comparable segment of the plot is 3 kbp-2 Mbp.

- Poor agreement was observed in the short-range contact region (up to 10 kbp), suggesting a mismatch between the chosen interaction potentials and the underlying physical parameters (see Methods).
- A generally reduced contact density was noted across the 3 kbp to 2 Mbp range.
- The fluctuations in the derivative of the contact probability are consistent with the stochastic noise typical of complex molecular systems under limited sampling.
- The Step potential produced results more consistent with the experimental data than the DPD potential, primarily on the basis of closer visual agreement of the simulated P(s) curves with the experimental profiles and superior performance according to quantitative metrics assessing differences between simulated and experimental contact probability distributions (see Methods).

Although both models using Step and DPD potentials generally reproduce the shape of the experimental P(s) curve, significant differences in detail, especially in the short-range range and the overall level of contact density, point to the limitations of the parameters and potentials used. These differences highlight the need for careful selection of physical parameters and normalization approaches to achieve better agreement between simulation and experiment.

### Quantitative Agreement Metrics are Sensitive to Normalization Point in Model Fits

In addition to direct visual comparison, we employed several quantitative metrics to assess the similarity between the simulated and experimental contact probability curves. However, a key methodological issue arises: P(s) curves are relative by nature and require normalization at a chosen genomic distance. For example, in (12), normalization was performed at 3 kbp, and comparisons were made over the range of 3 kbp to 2 Mbp. In contrast, (18) applies normalization at 10 kbp or, in some cases, 100 kbp to emphasize longer-range contacts.

These differences in normalization introduce significant variability in commonly used metrics, including the coefficient of determination (R^2^) for P(s), root mean square error (RMSE), mean absolute error (MAE), maximum error (ME), and average ratio. In contrast, R^2^ calculated for the derivative of P(s) remains unaffected by normalization since the derivatives are independent of the absolute scaling of P(s) (Fig 1B).

We observed that the R^2^, RMSE, and MAE values generally increase with normalization points approaching ∼100 kbp, reaching a maximum near 1 Mbp. The ME metric exhibits a similar trend, although its maximum occurs at approximately 100 kbp. Beyond these points, all the metrics based on P(s) begin to decline in quality, especially as the analysis approaches the 2 Mbp upper boundary. In contrast, the average ratio metric peaks at 20–30 kbp and shows a pronounced decrease beyond 1 Mbp.

This analysis was repeated across all simulation conditions presented in this study (see Appendix). For consistency, we selected 3 kbp as the normalization point for determining the best-fit parameters throughout the paper, aligning with the normalization strategy used in (12).

It is important to define and consistently apply a normalization point when analysing P(s) curves, as variations in this choice can substantially affect the values of similarity metrics and hinder direct comparisons across different studies. Moreover, in certain cases, the choice of normalization point can even influence which model appears to provide a better fit to the experimental data (S1-S2 Figs). Fixing the normalization point throughout the analysis enhances the robustness and reproducibility of model benchmarking. Unlike metrics based on the absolute values of P(s), this derivative-based R^2^ is independent of overall scaling and normalization, providing a more rigorous and unbiased assessment of the model’s ability to reproduce the shape of contact probability profiles. Furthermore, this metric quantifies how well the model approximates the experimental derivative relative to the average slope observed in the experimental data over the comparison range. Since the average slope of the P(s) derivative is directly linked to the underlying fractal dimension of the polymer backbone in the absence of extrusion-driven loops, the derivative-based R^2^ can be regarded as a physically grounded and informative measure of model fidelity.

### Dynamic Behavior of Condensin I is Inconsistent with Experimental Signatures

The use of static extruders with an exponential loop size distribution raises important questions regarding their validity for modeling the dynamic behavior of SMC complexes. In (12), static extruders are employed in conjunction with externally imposed large-scale helical constraints, which promote chromosome compaction. In contrast, (18) demonstrated that a similar helical structure can emerge spontaneously in simulations with dynamic extruders, even in the absence of external spatial constraints. In our simulations, however, we did not observe the formation of helical structures under either static or dynamic extrusion regimes (see Appendix), in contrast to the results reported by (18) for dynamic models.

A related consideration is the choice of parameters for modeling dynamic extruders (mitotic condensins). Loop size can be limited either by the intrinsic processivity of condensin or by natural roadblocks (for example, other extruders, cohesin-specific barriers such as CTCF, transcription, and related obstacles). Notably, condensin has been reported to bypass even large DNA-bound obstacles (21). To date, there are no well-established strong barriers to condensin motion along DNA, and transcriptional activity during mitosis is minimal (26). Within this context, the key parameter is the extruder lifetime (the mean residence time on DNA), which, together with the extruder loading density, directly determines the characteristic loop size. We estimated the baseline lifetime via the following relation:

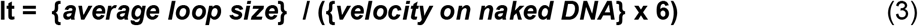

where the average loop size for condensin I is taken to be 100 kbp, the extrusion velocity is set to 1 kbp/sec (27), and the 1/6 factor accounts for the effective scaling of the polymer modeled as a chain of 10 nm nucleosome beads. Since the beads are 10 nm, which corresponds to 30 bp of naked DNA, and each nucleosome accounts for approximately 200 bp of naked DNA, and since the extruder makes spatial steps, the effective step distance is approximately 200/30 = ∼6 times greater than that of naked DNA, the velocity on chromatin is 6 times greater than that on naked DNA. This yields a baseline lifetime of approximately 17 seconds, which is sufficient for ∼63% of extruders to reach the next adjacent extruder along the polymer. In the simulations, each condensin complex has a probability of 1/lt to detach from the polymer at each time step and is immediately reloaded at a random location. SMC complexes perform this translocation through discrete spatial steps: recent single-molecule studies have demonstrated that condensin and cohesin translocate DNA in steps with a median size of ∼40 nm on naked DNA (28,29), with each step accompanied by a DNA twist of approximately −0.6 rotations (30). This mechanistic foundation (discrete spatial stepping coupled with DNA wrapping) supports our coarse-grained representation of extrusion as a sequence of bead-displacement events in our simulations.

To explore the effect of extruder dynamics, we simulated three different lifetimes:

- **17 seconds**, corresponding to the time required for condensin I to extrude its average loop size (∼100 kbp);
- **68 seconds**, corresponding to the time required for condensin II to extrude its average loop size (∼400 kbp);
- **136 seconds**, which was used as a control to investigate the effects of prolonged binding.

As shown in Fig 2, we observe marked differences in the contact density profiles between the dynamic and static extrusion regimes. For extruders with a 17-second lifetime, the contact density is elevated relative to the experimental data at distances in the 10–100 kbp range, followed by a steep decline. This reflects the short binding duration, during which many extruders fail to encounter obstacles before detaching; approximately one-third dissociate within 17 seconds.

**Fig 2.**
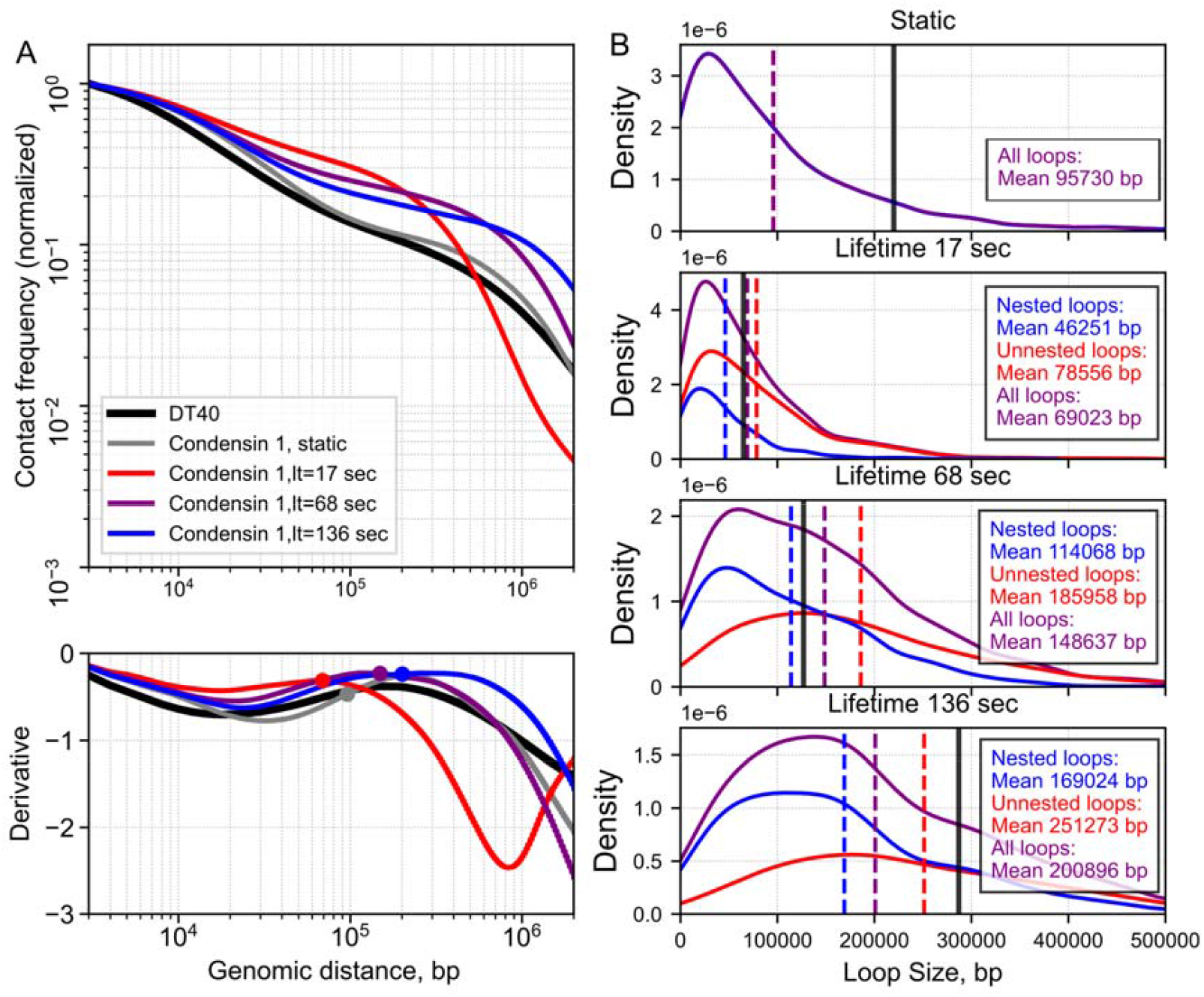
Effect of extruder lifetime on contact probability P(s) profiles. A) P(s) curves in the range of 3 kbp - 2 Mbp for experimental data for the chicken cell line DT40 (black line), aggregated model data. Static (gray line) and dynamic extruders with three lifetimes and base physical parameters (red, blue, purple lines). The large dots in the derivative graphs correspond to the mean loop size; B) Loop size distributions for different lifetimes. All the graphs are normalised on the area under the all-loops graph. The purple line represents all extruders, the red line represents unnested extruders, and the blue line represents nested extruders. 1) Static extruders, 2) Dynamic extruders lt = 17 sec, 3) Dynamic extruders lt = 68 sec, 4) Dynamic extruders lt = 136 sec. Black line - corresponding derivative peak size.

For the 68-second condition, the number of short-range contacts (<40 kbp) is reduced relative to the 17-second case because fewer short-lived extruders exist. However, the contact density increases significantly in the 50–200 kbp range, aligning more closely with the experimental data in this interval. Beyond this range, the model again overestimates contact decay.

The 136-second extruders exhibit similar behavior up to ∼20 kbp, but the contact density is slightly lower than that in the 68-second case up to ∼70 kbp. However, between 70 and 600 kbp, the contact probability profile closely matches the experimental curve. Beyond 1 Mbp, discrepancies reemerged across all conditions, which is consistent with the finite influence range of loop extrusion observed in the experimental system.

To further investigate the impact of extruder lifetimes on the three-dimensional organization of chromatin, we analysed the resulting distributions of loop sizes under different dynamic conditions. This analysis explicitly considers the formation of nested loops, which occur when new loops are extruded within preexisting loops. Importantly, that z-loop formation (21) does not occur in our model, as extruders are not permitted to pass through one another.

For extruders with a 17-second lifetime, the loop size distribution follows an approximately exponential profile (Fig 2B), closely resembling that observed for static extruders. In this regime, nonnested loops significantly outnumber nested loops. As the extruder lifetime increases, the average loop size increases accordingly. Because the total number of extruders on chromatin remains constant, the probability of new extruders binding within existing loops increases, leading to an increase in the number of nested loops. With further increases in lifetime (e.g., 136 seconds), the ratio of nested to nonnested loops continues to rise, indicating progressive structural complexity.

Previous studies have established a correlation between the average loop size and the position of the peak in the derivative of the P(s) curve (7,31,32). While our findings support the overall trend — namely, that larger average loop sizes are associated with a rightward shift of the derivative peak — we also observe quantitative discrepancies between these two measures. Specifically, under the 136-second lifetime condition (Fig 2), the position of the derivative peak substantially exceeds the corresponding average loop size.

A similar discrepancy is observed in the case of static extruders when Gaussian noise exceeds 50 nm (S1 Appendix). Although the loop size is fixed at 100 kbp, the derivative peak shifts from ∼90 kbp to nearly 300 kbp as the Gaussian noise increases (S1 Appendix). These findings suggest that while the derivative of P(s) is a useful qualitative indicator of loop size, it may be quantitatively unreliable under certain conditions, such as high noise levels or extensive nested loop formation. Accordingly, the position of the derivative peak should be treated only as a first-order approximation of the extruder-imposed loop size in the experimental data. When greater precision is needed, a targeted parameter sweep in the vicinity of the derivative peak is recommended to refine the estimate.

As shown in Fig 2, increasing the extruder lifetime leads to a marked rise in the number of nested loops. This effect is associated with a reduction in the effective number of active extruders, resulting from spatial overlap and the hierarchical nature of nested loop organization. To address this limitation, we implemented an alternative strategy that introduced spatial exclusion rules, whereby condensin complexes are restricted to loading only within interloop regions, and not within existing loops (Fig 3).

**Fig 3.**
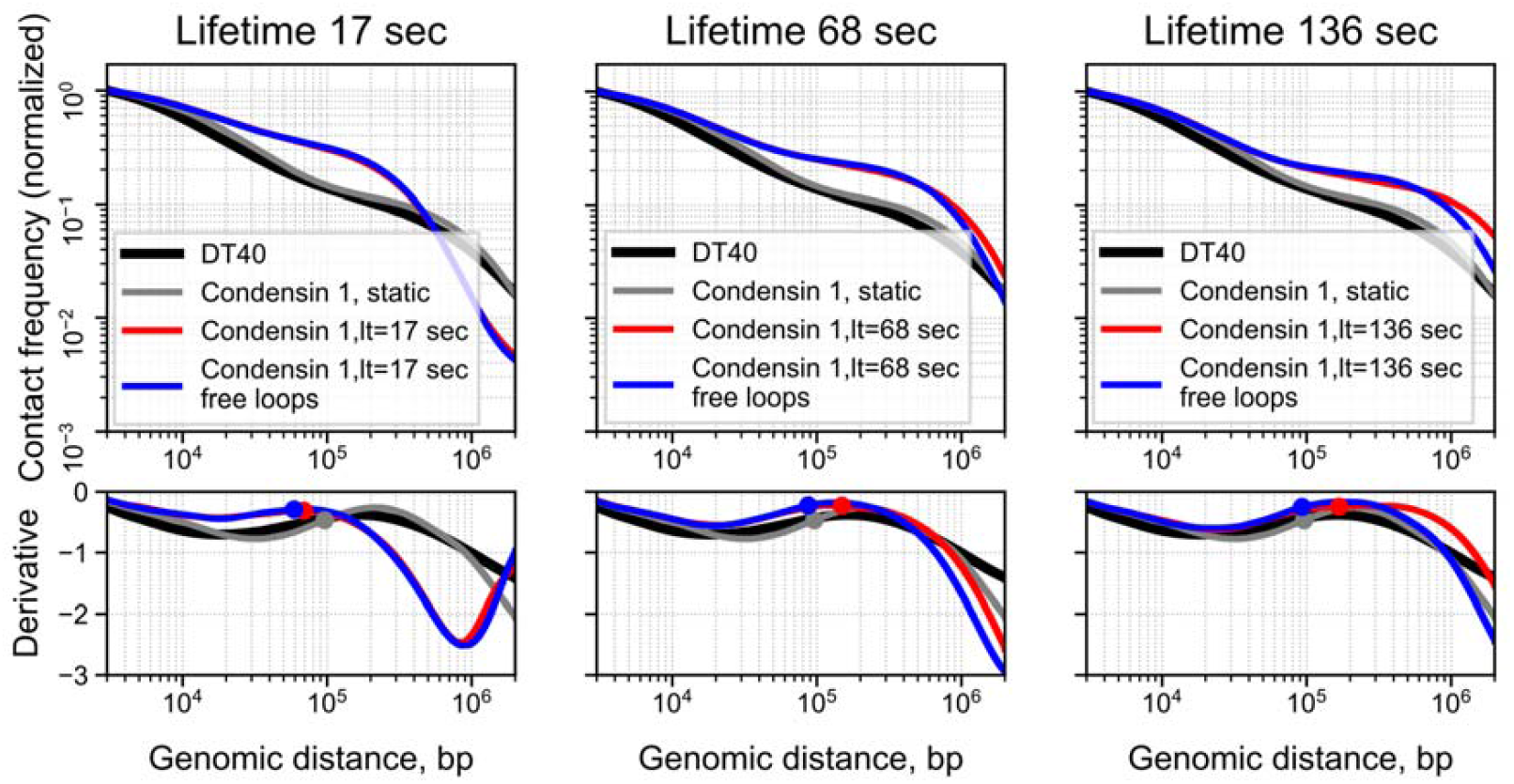
Impact of constrained extruder dynamics on contact probability P(s) profiles across lifetimes. P(s) curves in the range of 3 kbp - 2 Mbp for experimental data for the chicken cells line DT40 (black line) and aggregated model data. The large dots in the derivative graphs correspond to the mean loop size.

Under this modified framework, the loop size distribution for extruders with a 17-second lifetime remained unchanged, in agreement with theoretical expectations. However, at longer lifetimes, the distributions shifted substantially, becoming more similar to those observed in the static extruder configuration, as reflected in both the P(s) curves (Fig 3) and loop size distributions (S3 Fig).

The rationale for this exclusion model stems from experimental observations that condensins can induce chromatin twisting or torsional stress (30), which may affect the physical properties of the substrate and reduce the likelihood of additional extruders binding to already extruded loops. It is therefore theoretically plausible that torsional tension contributes to a reduced probability of condensin loading in regions already occupied by other extruders.

None of the tested models outperformed the static extruder model in terms of quantitative agreement with the experimental data. Specifically, the average ratio metric for the static model with optimal Gaussian noise (σ = 200 nm, see Methods) was 1.16. For the DPD potential model with its best-performing noise level (σ = 100 nm), the value was 1.23 (S1 Appendix). The dynamic extruder model with its optimal lifetime (136 sec) yielded a value of 1.37 (S1 Fig), and the dynamic model with restricted loading (i.e., extruders prohibited from binding within existing loops) achieved a value of 1.35 (S2 Fig) under the optimal lifetime (136 sec).

A similar trend was observed for the derivative R^2^ metric, which reflects the agreement in slope between the modeled and experimental P(s) curves. The static model with σ = 200 nm achieved a derivative R^2^ value of 0.98; the DPD model with σ = 100 nm produced 0.97; the unrestricted dynamic extruder model with the optimal lifetime (136 sec) yielded 0.72; and the restricted dynamic model yielded 0.77 under the optimal lifetime (136 sec).

## Discussion and conclusions

In the present study, we systematically compared static and dynamic loop extrusion models for modeling mitotic chromosome folding via chicken Hi-C data. Our work aimed to clarify whether these two approaches are interchangeable or whether they lead to fundamentally different results.

We showed that although both models can be tuned to approximate the general shape of the contact probability curve P(s), the static extruder model demonstrated the best quantitative agreement with the experimental data compared with all the tested dynamic models. In particular, the static model with an optimally chosen noise level (σ = 200 nm) and interaction potential Step achieved the best performance on metrics such as the average ratio and coefficient of determination (R^2^) for the derivative of P(s). However, given the breadth of the hyperparameter space (including extruder abundance, translocation velocity, and lifetime; bead–bead interaction energies; the presence or absence of PBS boxes; and the level of Gaussian smoothing (σ)), it remains plausible that an alternative interaction potential could yield a superior approximation to the experimental data under a different, yet biologically reasonable, parameterization.

Our results highlight key limitations of dynamic models in this context. The increase in extruder lifetime required to reproduce contacts at large genomic distances resulted in the formation of an excessive number of nested loops. This, in turn, distorted both the loop size distribution and the P(s) curve. In addition, we found that under high noise conditions or with many of nested loops, the position of the peak of the P(s) derivative ceases to be a reliable indicator of the average loop size.

We qualitatively and quantitatively compared these two modeling paradigms within a single platform (Polychrom), which allowed us to identify their intrinsic differences. We also demonstrated a strong dependence of the results on the choice of parameters such as the normalization point of the P(s) curves and the level of spatial noise, and proposed using the R^2^ of the P(s) derivative as a more reliable metric for comparing the models.

The novelty of our study is that we perform, for the first time, a direct comparison of static and dynamic models of extrusion within a single software framework using parameters from a recent work.

Thus, our study confirms that models with static extruders, despite their apparent simplicity, remain powerful and adequate tools for describing the final structure of mitotic chromosomes. Across the conditions examined, the static model of the mitotic chromosome provided the closest agreement with the experimental data. In contrast, dynamic extrusion models in which extruders stall upon collision consistently yield poorer fits. Attaining comparable performance with a dynamic framework requires extruder lifetimes on the order of the mitotic phase, which effectively produces loop size distributions indistinguishable from those of the static model. Even with an added spatial-exclusion rule that prohibits loading within preexisting loops, satisfactory agreement still necessitates long lifetimes, again rendering the behavior functionally equivalent to the static configuration.

An alternative model assumes that condensins bind DNA once at mitotic entry, extrude loops until they encounter other complexes, and remain stably associated until the end of mitosis. However, this model would require the introduction of additional mechanisms. For example, the experimentally observed nonzero gaps between loops (12) could be explained by stalling forces arising from chromatin tension exceeding ∼2 pN. Similarly, the progressive compaction of chromosomes and the upwards shift in the second diagonal of contact maps, despite relatively stable loop sizes for condensin II (12), may reflect the influence of loop extrusion– independent mechanisms, such as the activity of topoisomerase II (10).

## Methods

### Polymer Models

We constructed polymer models of chromosomes via polymer dynamics simulations. The simulations were performed via a package based on open2c/polychrom (Imakaev et al., 2019), which integrates parameters from both static and dynamic simulations as described in Samejima et al. (2025).

The simulation workflow consists of two main stages:

1. One-dimensional (1D) simulation The polymer chain is represented as a 1D lattice composed of beads, each 10 nm in size and corresponding to 200 base pairs (bp).
  1.1. Dynamic extruders, representing condensin I complexes, are modeled as walker objects with two legs that move probabilistically in opposite directions. At each time step, only one leg is active and attempts to move up to approximately 50 nm (5 beads) in its respective direction. The active leg switches with a 50% probability, effectively simulating two-sided loop extrusion. The positions of the legs define the loop boundaries extruded by condensins.
  1.2. Condensins bind randomly to pairs of adjacent free lattice sites and extrude loops until unbinding occurs, after which they immediately rebind at random locations. When two legs of different extruders collide, extrusion stalls until one leg unbinds.
  1.3 Static extruders are also modeled as walker objects, but with legs loaded onto lattice sites following an exponential distribution with a mean loop size of 100 kbp. These static extruders do not perform extrusion or unbinding.
  1.4. The condensins identified in the 1D simulation are subsequently implemented as additional harmonic constraints in the 3D polymer model.
  1.5. The simulation time step is 0.1 seconds. The condensin I velocity is set to 1 kbp/s on naked DNA (27), corresponding to a step size of 50 nm (28); this translates to approximately 6 kbp/s on condensed chromatin. The total simulation duration is 1800 seconds.
  1.6 All 1D modeling was performed via our scripts (https://github.com/Three-point/1D-polymer-models)
2. Three-dimensional (3D) simulation This stage employs molecular dynamics simulations using normalized units, where particles have a mass of 1 unit and a bead radius of 0.5 units, with one length unit corresponding to 10 nm.
  2.1. The monomers are connected by harmonic springs (polychrom.forces.harmonic_bonds) with a bond length of 1 unit and a wiggle distance of 0.1 units.
  2.2. The polymer stiffness (polychrom.forces.angle_force) is modeled with a persistence length of 150 bp, equivalent to three nucleosome linkers of 50 bp each, implemented via an angular force with k=3.
  2.3. Repulsive forces prevent spatial overlap via polynomial repulsion (polychrom.forces.polynomial_repulsive) with Erep = 3.0, mimicking real polymer behavior (Formula 1). The collision rate is set to 0.01 inverse picoseconds, and the Langevin integrator is used with an error tolerance of 0.01. Each time step comprises 1000 molecular dynamics steps.
  2.4. No external spatial constraints are applied to the polymer in this work. This differs from (12), where cylindrical and pulling constraints were used to achieve the required densities and patterns.

Each dataset comprises five independent 40 Mbp chains, each initialized with distinct condensin positions and starting conformations generated by random walks.

Two different potential energies were used for nonbonded bead repulsion.

- “Step potential” (22):
- “DPD potential” (23):

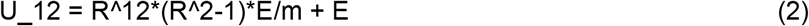

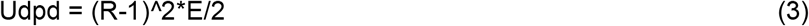

where R is the distance between beads in dimensionless units, E is the repulsion energy coefficient, and m is the normalisation constant for this degree of equation.

### Experiment types

There 11 types of model parameters were used:

- Static extruders. The positions of the extruders placed on the lattice with an exponential distribution of loops once do not change with time (no move, binding/unbinding):
  - Step potential
  - DPD potential (Groot and Warren, 1997, Formula 2), where Erep = 3.0 kT, k = 0 (polymer stiffness), and the other parameters are equal to the Step potential.
- Dynamic extruders. All the physical parameters are taken from the Step potential. They unbind from the lattice with a probability inversely proportional to the lifetime and then instantly bind back at random site. Then, they make one step by one of the legs chosen randomly (50% chance to switch direction) on 5 beads each time step with a probability sufficient for the target extrusion speed:
  - Lifetime = 17 sec for each extruder. This is enough for ∼60% of the extruders to extrude 100 kbp (if no obstacles are encountered), forming a distribution closest to exponential. There are low numbers of nested loops (loops on other loops) and a dominating number of unnested loops.
  - Lifetime = 68 sec for each extruder. Nested loops started dominating in numbers at short distances.
  - Lifetime = 136 sec for each extruder. Nested loops are dominant in terms of number for all loop sizes.
- Dynamic extruders that are unable to load at already existing loops, they load only at gaps between loops.
  - Lifetime = 17 sec for each extruder.
  - Lifetime = 68 sec for each extruder.
  - Lifetime = 136 sec for each extruder.

### Generation of Hi-C Maps

For each polymer chain in the simulations, we extracted conformations represented by the spatial coordinates of beads at the following time points: 1080 s, 1440 s, and 1800 s. This selection is based on the assumption that a 360 s interval (exceeding three condensin lifetimes) suffices to establish a new loop distribution under dynamic conditions.

Each bead position within a conformation was subsequently perturbed by adding a random Gaussian displacement with standard deviation σ along each spatial axis. By repeating this perturbation process, we generated 100 new conformations from each original conformation, resulting in a total of 5×3×100=1500 conformations per dataset.

For each conformation, pairs of 200 bp beads located within 11 nm (contact radius, 1.1 of the bead radius) were identified as contacts, from which a contact map in .mcool format at 200 bp resolution was constructed (https://github.com/Three-point/1D-polymer-models). Unless otherwise specified, the default value of σ was set to 200 nm, as this value yielded the best agreement with the experimental P(s) curve for static extruder conditions (see S1 Appendix).

Processed Hi-C data (.mcool) from the DT40 cell line of Gallus gallus (https://www.ncbi.nlm.nih.gov/geo/query/acc.cgi?acc=GSE262525), sample GSM8171168, were used as experimental data.

The P(s) curves were computed via the cooltools (version 0.5.1) and cooler (version 0.9.1) packages with curve Gaussian smoothing (σ=0.12) and excluding the first two diagonals of the Hi-C contact matrix.

### Quantitative Comparison of P(s) Curves

To quantitatively compare the contact probability curves P(s), we focused on the genomic distance range from 3 kbp to 2 Mbp, which encompasses the effective range of condensin I activity (typical loop size ∼100 kbp). Prior to metric evaluation, all curves were normalized at a selected genomic coordinate within this range. Given the logarithmic scaling of the genomic distance axis, discrepancies at larger distances exert a disproportionately strong influence on agreement metrics. Therefore, the analysed range was subdivided into 100 uniformly spaced intervals on a log scale.

The following metrics were used to assess the agreement between experimental and simulated P(s) curves in log10–log10 space:

- **Coefficient of determination (R**^**2**^): Values approaching 1 indicate strong agreement; negative values suggest that the model performs worse than a horizontal mean line.
- **Root Mean Square Error (RMSE):** Lower values denote better fit and reduced variance from the experimental curve.
- **Mean Absolute Error (MAE):** Measures the average magnitude of differences; lower values indicate better agreement.
- **Maximum Error (ME):** Captures the largest single deviation between curves; lower values are preferred.
- **Average Ratio:** At each point, the modeled P(s) value is divided by the corresponding experimental value. If the ratio is <1, its reciprocal is taken to ensure symmetry. The final value is the arithmetic mean of these normalized quotients across all distances. A value close to 1 reflects excellent fit, whereas larger values indicate greater overall deviation.

To compare the derivatives of the P(s) curves, only the **R**^**2**^ **metric** was employed. This choice is justified by its invariance to normalization, as it captures the similarity in slope and shape of the curves, independent of absolute scaling.

### Loop Distribution Analysis

Loop positions were extracted at identical time points (1080 s, 1440 s, and 1800 s) for each simulated cell and categorized into the following groups:

- **Nested loops**: loops fully enclosed within the genomic span of another loop.
- **Unnested loops**: loops that are not nested within any other loop.
- **All loops**: the complete set of loops detected, regardless of nesting status.

For each extruder, the genomic span—defined as the distance between its two extrusion legs in base pairs was calculated. The resulting distributions of loop sizes for each category were visualized via Kernel Density Estimation (KDE) plots, generated with the Seaborn Python library.

## Funding

The degron analysis in this work was supported by the grant of the state program of the «Sirius» Federal Territory «Scientific and technological development of the «Sirius» Federal Territory» (Agreement №26-03, 27/09/2024) (A.S.). Modelling part of the study was supported by the Ministry of Science and Higher Education of the Russian Federation, grant no. FSUS-2024-0018 (T.L.). The funders had no role in the study design, data collection and analysis, decision to publish, or preparation of the manuscript.

## Acknowledgements

Computational resources were provided by HPC facilities at the collaborative center «Bioinformatics» of ICG SB RAS (funded by the Ministry of Education and Science of the Russian Federation, state project FWNR-2022-0019). The original manuscript text was composed by authors; proofreading was conducted with the assistance of ChatGPT version 5, after which the text was corrected and edited by the authors.

## Author contributions

T.L. conceptualized the study. Bioinformatic analyses and polymer simulations were carried out by V.K., with contributions from T.L. and A.S. V.K. drafted the manuscript with contributions by T.L. and A.S. All authors reviewed, revised, and approved the final manuscript.

## Ethics approval and consent to participate

Not applicable

## Consent for publication

Not applicable

## Data Availability Statement

All the data and code underlying the findings are fully available. The simulation code is deposited at https://github.com/Three-point/Chrom_polymer_models. The simulation outputs (contact maps, .csv files and configuration files) are deposited at Zenodo (DOI: 10.5281/zenodo.17828395). Figure source data are provided in the Supporting Information.

## Competing interests

The authors declare that they have no competing interests

## Supporting information

**S1 Fig.**
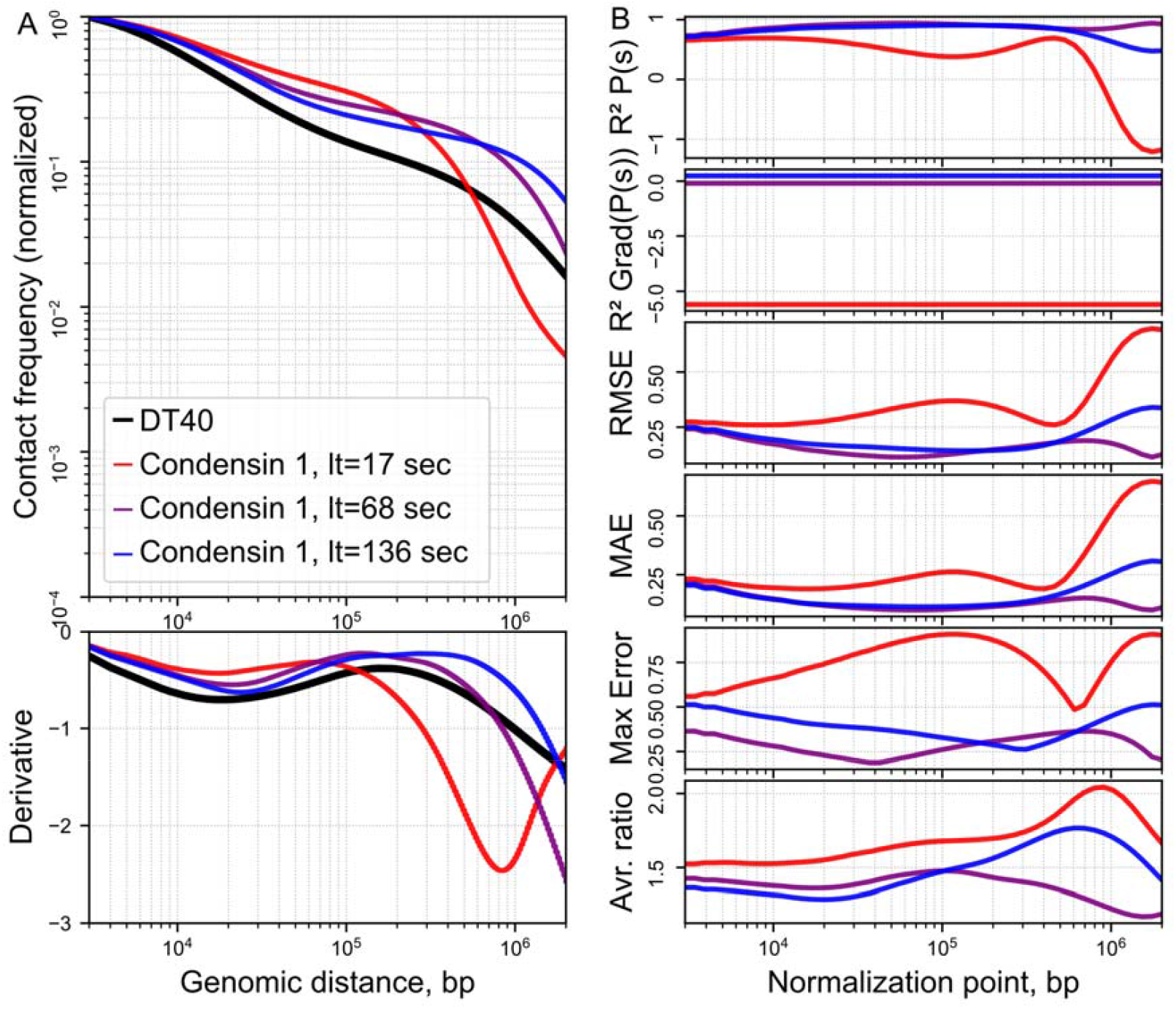
Effect of extruder lifetime on contact probability P(s) profiles with different metrics. A) P(s) curves in the range of 3 kbp - 2 Mbp for experimental data for the chicken cell line DT40 (black line) and aggregated model data. Static (gray line) and dynamic extruders with three lifetimes and base physical parameters (red, blue, and purple lines). The large dots in the derivative graphs correspond to the mean loop size; B) Results of comparing the segment experiment DT40 P(s) with the model data for the dynamic extruders over three lifetimes with shifting normalisation points on different metrics. The normalization point shifts within the range of 3 kbp to 2 Mbp. The comparable segment of the plot is 3 kbp-2 Mbp. The sigma-noise is 200 nm.

**S2 Fig.**
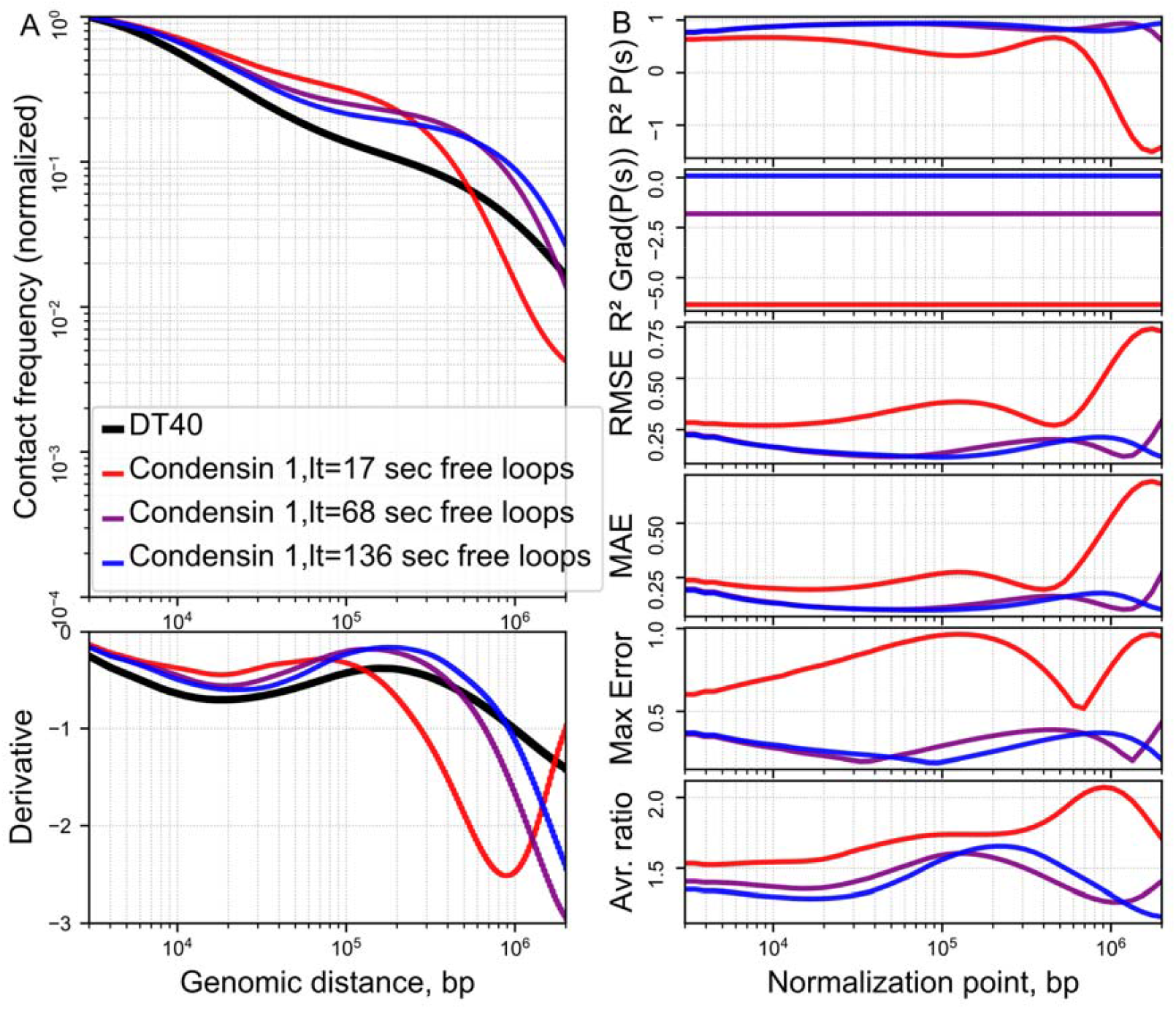
Effects of extruder lifetime free loops on contact probability P(s) profiles with different metrics. A) P(s) curves in the range of 3 kbp - 2 Mbp for experimental data for the chicken cell line DT40 (black line), aggregated model data. Static (gray line) and dynamic extruders, which are unable to load at existing loops, on three lifetimes with base physical parameters (red, blue, purple lines). The large dots on the derivative graphs correspond to the mean loop size; B) Results of comparing the segments experiment DT40 P(s) with the model data on different metrics. The normalization point shifts within the range of 3 kbp to 2 Mbp. The comparable segment of the plot is 3 kbp-2 Mbp. The sigma-noise is 200 nm.

**S3 Fig.**
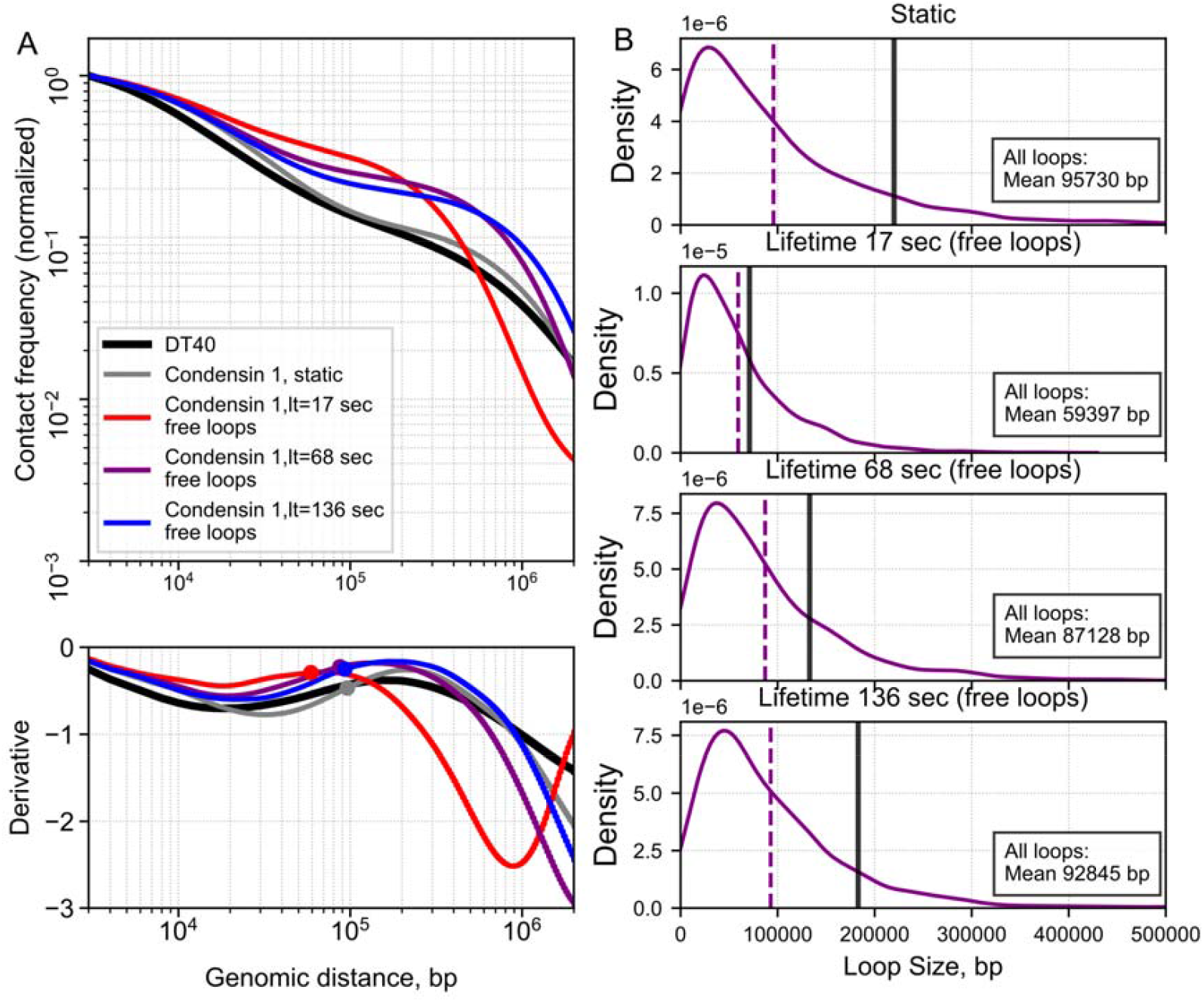
Effect of extruder lifetime free loops on contact probability P(s) profiles with loop size distributions. A) P(s) curves in the range of 3 kbp - 2 Mbp for experimental data for the chicken cell line DT40 (black line), aggregated model data. Static (gray line) and dynamic extruders, which are unable to load at existing loops, on three lifetimes for Step potential parameters (red, blue, purple lines). The large dots in the derivative graphs correspond to the mean loop size; B) Loop size distributions for different lifetimes. All the graphs are normalised on the area under the all-loops graph. Purple line - all extruders. 1) Static extruders, 2) Dynamic extruders lt = 17 sec, 3) Dynamic extruders lt - 68 sec, 4) Dynamic extruders 136 sec.

**S1 Appendix. Artificial Noise Effects and Their Role in Model-Data Reconciliation; Analysis of Spiralization in Chromosome Models Using C(s) Correlation**.

## Notes

### Competing Interest Statement

The authors have declared no competing interest.

### Summary of Updates

Image readability has been improved. The introduction has been expanded, several sections have been renamed for better clarity, and the text has been comprehensively rewritten to be more accessible with refined language. Additionally, one section has been relocated to the appendix, and a new supplementary section has been added to the appendix

