## Supplementary material for "Effects of Extruder Dynamics and Noise on Simulated Chromatin Contact Probability Curves": Si Appendix

### **Artificial Noise Effects and Their Role in Model-Data Reconciliation**

Fig 1 illustrates that contact-based analyses under conditions of sparse interaction sampling are inherently sensitive to stochastic fluctuations, necessitating cautious interpretation of contact density variations. One commonly used strategy to address this limitation is the introduction of positional noise, which can reconcile discrepancies between simulated and experimental data by increasing the effective number of contacts. In (Samejima et al., 2025), this is achieved by applying Gaussian displacements to the 3D coordinates of polymer conformations along all spatial axes. This approach effectively generates ‘additional’ conformations, thereby increasing the number of distant, probabilistic contacts—albeit at the cost of uniform short-range contact frequency. It is important to note that while increasing the magnitude of noise (σ) smooths the contact probability curve, excessive noise can obscure or eliminate the underlying structural features of chromatin.

The rationale behind this noise addition is rooted in the observed noisiness of Hi-C data, which often displays an almost uniform distribution of short-range contacts. In contrast, simulations generally lack such uniformity, leading to an underestimation of short-distance contact frequencies. Notably, switching the restriction enzyme from DpnII (used in standard Hi-C) to MNase (used in Micro-C) introduces more frequent cutting (~200 bp for MNase vs. ~1–5 kbp for DpnII), yielding a less uniform distribution of short-range contacts and better resolution of chromatin structure (Akgol Oksuz et al., 2021). In human H1-hESC cells, the most pronounced changes in the derivative of the contact probability occur within the ~1 kb to ~100 kb range, which corresponds closely to the characteristic size of interphase chromatin loops (~100 kb).

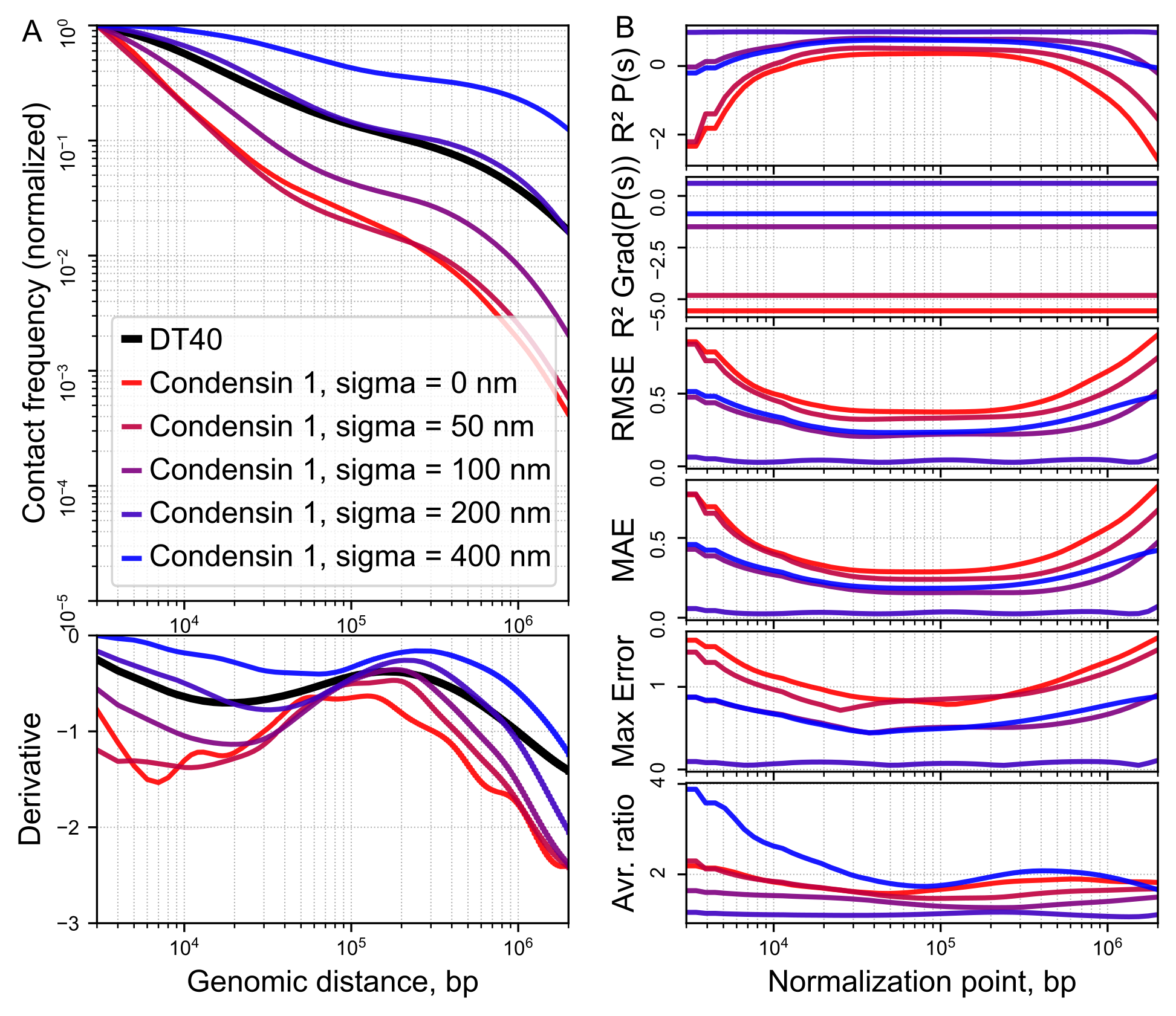

**Fig A1. Effects of spatial displacement noise value with “Step potential” on contact probability P(s) profiles.**
A) Smoothing of P(s) model curves with different sigma-noises in the range of 3 kbp - 2 Mbp for experimental data for the chicken cell line DT40 (Samejima et al., 2025) (black line), aggregated model data for the Step potential; B) Results of comparing segment experiments P(s) with the model data with shifting normalisation point on different metrics. The normalization point shifts within the range of 3 kbp to 2 Mbp. The comparable segment of the plot is 3 kbp-2 Mbp. The sigma noise is 200 nm.

When σ was varied over a range of [50, 100, 200, 400] nm (Fig A1A), a predictable decrease in contact density was observed, primarily due to the loss of short-range contacts below 10 kbp. The normalization point for these analyses was fixed at 3 kbp. Notably, the normalization had no impact on the derivative of P(s), which revealed clear divergence between high-noise model curves and experimental data.

We observed that increasing σ shifted characteristic local maxima and minima in the P(s) derivative toward larger genomic distances. This is consistent with the idea that smaller loops are more susceptible to noise-induced perturbation. A comparable analysis performed via the DPD potential (Fig A2) yielded qualitatively similar results.

In (Samejima et al., 2025), the optimal σ was reported as 49 nm, producing an average ratio metric of approximately 1.05 via the DPD potential. In our Step potential implementation, we observed a similar value (~1.16, Fig A1B) at σ = 200 nm, while the DPD-based implementation yielded a best value of ~1.23 at σ = 100 nm (Appendix). These σ values are consistent with the DpnII digestion scale, where 49 nm corresponds to roughly 5 nucleosomes (~1,000 bp), and 200 nm to ~20 nucleosomes (~4,000 bp).

Based on this analysis, we selected the Step potential with σ = 200 nm as the default configuration for all subsequent simulations, due to its favorable performance across multiple fit metrics.

Addition of positional noise plays an important role in smoothing the contact probability curves and improving the fit of the model to experimental data by increasing the effective number of contacts. However, excessive increase in the noise level leads to the loss of structural features of chromatin, which limits the accuracy of interpretation of the results. Thus, the optimal level of Gaussian noise must be carefully selected, taking into account the scale of restriction enzyme cuts and the characteristic sizes of chromatin loops, to ensure a balance between the realism of the modeling and the preservation of biologically significant structures.

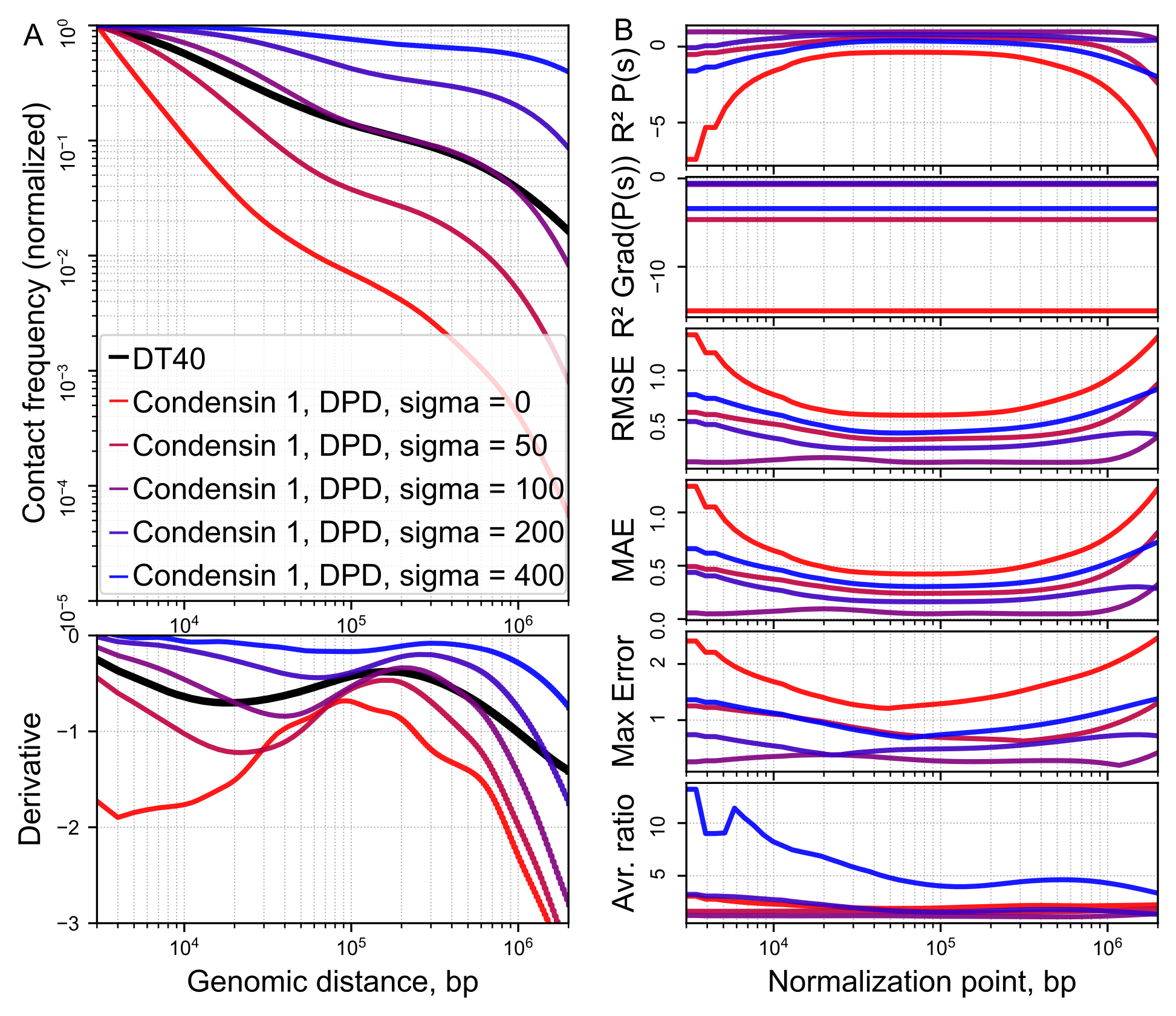

**Fig A2. Effects of spatial displacement noise value with “DPD potential” on contact probability P(s) profiles.**
A) Smoothing of P(s) model curves with different sigma-noises in the range of 3 kbp - 2 Mbp for experimental data for the chicken cell line DT40 (black line), aggregated model data for the DPD potential; B) Results of comparing segments experiment P(s) vs model data with shifting normalisation point on different metrics. The normalization point shifts within the range of 3 kbp to 2 Mbp. The comparable segment of the plot is 3 kbp-2 Mbp. The sigma noise is 200 nm.

### **Analysis of Spiralization in Chromosome Models Via C(s) Correlation**

To quantify the extent of spiralization (large-scale helical order) in our simulations and in those reported previously (Samejima et al., 2025), we computed the spatial correlation function C(s) (Dey et al., 2023, Fig A3).

The function C(s) measures the orientational memory along the chromatin fiber at a separation s. For a given curve parameterized as a sequence of 3D positions, we define tangent (bond) vectors as *v_i = r_(i+d) − r_i*, where *d* is the bead number shift. The vectors are normalized to unit length, and C(s) is computed as the averaged scalar product between all pairs of normalized vectors separated by s along the contour:
C(s)=⟨v_i ⋅ v_(i+s)⟩

where the average is taken over all possible i. The function oscillates if the structure contains periodic or helical (spiral) order, with the period corresponding to the helical turn. For a completely random coil, C(s) decays rapidly to zero, while for an ideal spiral, pronounced oscillations persist over long distances.

In chromosomal studies, spiralization is also often inferred from the appearance of a peak in the contact probability curve P(s) at a characteristic genomic separation. However, it is important to note that such a peak is only a suggestive indicator—various global or local folding mechanisms can produce similar features in P(s) without genuine spiral order.

In our simulated chromosome models, no clear peak in P(s) is observed, likely due to both the limited polymer length and the absence of constraints (e.g., geometric confinement, higher-order folding cues) that stabilize large-scale helicity. Correspondingly, the C(s) curves computed from our simulations exhibit only weak or absent periodicity, except potentially at the largest vector separation step d.

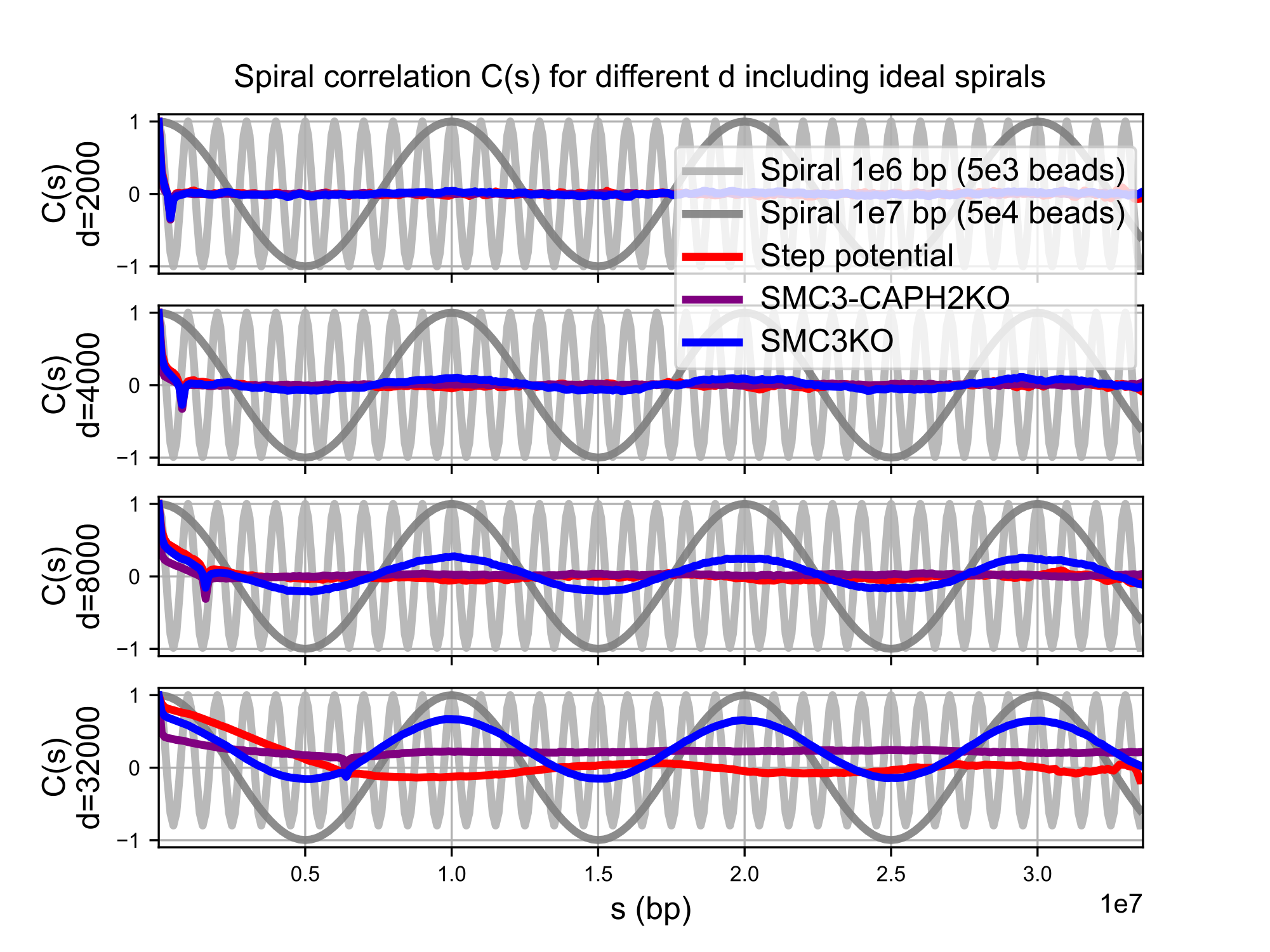

**Fig A3. Effect of the bead number shift on Spiral correlation C(s) profiles.**
Spiral correlation C(s) for different vector step sizes d including ideal spirals. Each subplot shows C(s) as a function of genomic distance s (in bp) for a fixed d. Gray lines: Analytical result for ideal spirals with pitches 1×10^6 and 1×10^7 bp, serving as periodic references. Red lines: Averaged C(s) for simulated model chromosomes. Purple/Blue lines: Experimental data for SMC3-CAPH2KO and SMC3KO (Samejima et al., 2025).

At small and moderate d, both simulated and most experimental traces show rapid loss of correlation with minimal periodicity, ruling out strong spiralization. Only in SMC3KO (Samejima et al., 2025) does the expected oscillation appear, revealing helical order in this perturbation.
